# Tightly-coupled inhibitory and excitatory functional networks in the early visual cortex

**DOI:** 10.1101/2021.08.27.457908

**Authors:** Haleigh N. Mulholland, Bettina Hein, Matthias Kaschube, Gordon B. Smith

## Abstract

Intracortical inhibition plays a critical role in shaping activity patterns in the mature cortex. However, little is known about the structure of inhibition in early development prior to the onset of sensory experience, a time when spontaneous activity exhibits long-range correlations predictive of mature functional networks. Here, using calcium imaging of GABAergic neurons in the early ferret visual cortex, we show that spontaneous activity in inhibitory neurons is already highly organized into distributed modular networks before visual experience. Inhibitory neurons exhibit spatially modular activity with long-range correlations and precise local organization that is in quantitative agreement with excitatory networks. Furthermore, excitatory and inhibitory networks are strongly co-aligned at both millimeter and cellular scales. These results demonstrate a remarkable degree of organization in inhibitory networks early in the developing cortex, providing support for computational models of self-organizing networks and suggesting a mechanism for the emergence of distributed functional networks during development.

## Introduction

Inhibition is crucial for shaping neural activity and response properties in mature cortex. GABAergic interneurons have been implicated in a range of computations, including response gain, stimulus discrimination, and network stabilization (for reviews, see Isaacson and Scanziani, 2011; Denève and Machens, 2016; Ferguson and Cardin, 2020). In the columnar visual cortex, inhibitory neurons actively shape response selectivity (Wilson, Scholl and Fitzpatrick, 2018) and are organized into functionally-specific networks with excitatory neurons (Wilson *et al*., 2017). However, relatively little is known about inhibition in the early cortex prior to the onset of sensory experience. GABAergic inhibition is initially absent in early development before progressively strengthening as the cortex matures (reviewed in Ben-Ari, 2002; Huang et al., 2007), raising the possibility that inhibition plays only a minor role in shaping early patterns of cortical activity.

Recent work in the ferret visual cortex demonstrated that prior to the onset of visual experience, excitatory activity is already highly structured, showing modular and distributed activity with long-range correlations (Smith *et al*., 2018), reminiscent of the columnar stimulus-evoked activity found in the mature cortex (Hubel and Wiesel, 1968; Blasdel and Salama, 1986; Weliky, Bosking and Fitzpatrick, 1996; Issa, Trepel and Stryker, 2000; Kara and Boyd, 2009; Smith, Whitney and Fitzpatrick, 2015). These early correlated networks are not abolished when silencing feedforward activity and predict future visually-evoked responses (Smith *et al*., 2018), suggesting a key role for early spontaneous activity in the development of mature cortical networks. However, the structure of inhibitory activity at this early stage of development remains unknown. Several distinct possibilities exist (Figure 1): Inhibition could be weak or absent in the early cortex, or be broad and unstructured, operating over a much larger spatial scale than excitation. In contrast, early inhibition might be organized and modular, but poorly aligned with excitation. Finally, the highly organized and co-aligned patterns of excitation and inhibition found in mature animals (Wilson *et al*., 2017) might already be present in the early cortex. Resolving this question is critical to both constrain models of early network development—many of which rely on structured intracortical inhibition—and to understand the degree to which inhibition contributes to early distributed patterns of cortical activity.

**Figure 1.**
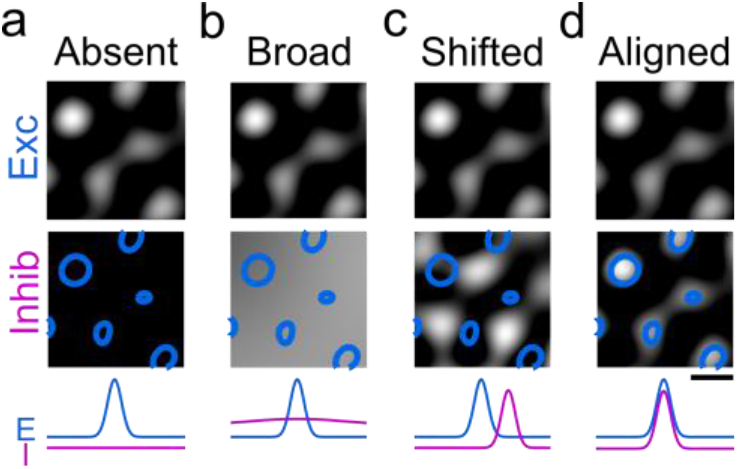
Possible modes of inhibition in the early cortex. Schematic showing potential arrangements of inhibitory and excitatory activity patterns in early cortex. *Top row:* Excitation is known to be highly modular and spatially structured. *Middle row:* Inhibition could be absent (a), broadly unstructured (b), or modular (c and d); if modular, the spatial arrangement of those modules could be shifted with respect to excitatory activity (c), or already well aligned as found in the mature cortex (d). Scale bar 1 mm.

Here we address this by using wide-field and two-photon calcium imaging of spontaneous activity to examine the structure of inhibitory networks in the developing ferret visual cortex. We first demonstrate *in vivo* that already in the early cortex, GABAergic signaling exerts a strong inhibitory effect. Next, by employing the inhibitory neuron-specific enhancer *mDlx* (Dimidschstein *et al*., 2016) paired with the genetically encoded calcium indicator GCaMP6, we find that inhibitory neurons exhibit modular patterns of activity that extend over several millimeters in the early visual cortex. These patterns reveal long-range correlated networks with precise local organization, highly similar in structure to those of excitatory networks. Furthermore, we find that inhibitory and excitatory networks show a remarkable degree of spatial consistency and co-alignment on both global and cellular scales. These findings clearly demonstrate the long-range and fine-scale organization of intracortical inhibition in the developing cortex, and support the ability of large-scale cortical networks to self-organize through precisely correlated local excitatory and inhibitory activity.

## Results

### Early spontaneous activity in inhibitory populations is modular and forms large-scale correlated networks

Over the course of development, GABAergic signaling switches from exerting depolarizing effects to hyperpolarizing due to the maturation intracellular chloride concentrations (Ben-Ari, 2002). In the ferret, functional GABAergic synapses have been shown to be present as early as postnatal day 20 (P20) (Dalva, 2010), but because these experiments utilized whole-cell recordings in cortical slices, it remains unclear whether these synapses exert an inhibitory effect at this age. To address this, we expressed GCaMP6s in excitatory neurons at P21-23 via AAV1-hSyn-GCaMP6s (Wilson *et al*., 2017) and imaged spontaneous activity prior to and following direct application of the GABA(A) antagonist bicuculline methiodide (BMI) to the cortex (Figure 2a-b). BMI resulted in a pronounced increase in the frequency (Figure 2c. p<0.001; baseline= 0.030 (0.016-0.050) Hz; BMI=0.200 (0.129-0.271) Hz, n=4 animals; Median, inter-quartile range (IQR), Wilcoxon rank sum test) and amplitude of spontaneous events (Figure 2d. p<0.001; Baseline=0.374 (0.190-0.535) ΔF/F, n (across 4 animals)=274 events; BMI=0.974 (0.470-1.220) ΔF/F, n=732 events; Median, IQR, Wilcoxon rank sum test), demonstrating that *in vivo* GABA exerts a strong net inhibitory action at this point of development.

**Figure 2.**
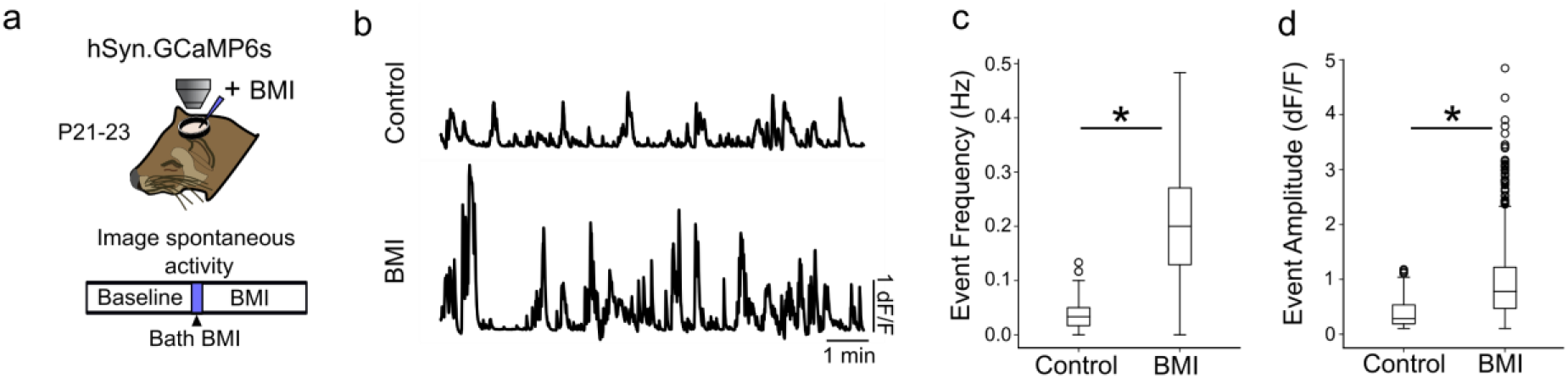
GABAergic signaling has net inhibitory effect on cortical activity by P21. **a.** Experimental schematic; Ferret kits were injected with syn.GCaMP6s at approximately P10, and spontaneous activity was imaged at P21-23 (n=4 animals) both before and after bath application of the GABA antagonist bicuculine (BMI). **b.** Average trace across imaging FOV of baseline spontaneous activity (*top*) and activity after BMI application (bottom) shows clear disinhibition of cortical activity. **c.** Event frequency after BMI application is significantly higher than baseline (p<0.001; baseline= 0.030 (0.0160.050) Hz; BMI=0.200 (0.129-0.271) Hz; Median, inter-quartile range (IQR), Wilcoxon rank sum test). **d**. Average event amplitude of BMI events are significantly higher than baseline (p<0.001; Baseline=0.374 (0.190-0.535) ΔF/F, n (across 4 animals)=274 events; BMI=0.974 (0.470-1.220) ΔF/F, n=732 events; Median, IQR, Wilcoxon rank sum test).

To examine the structure of inhibitory networks in early development, we then labeled GABAergic neurons via virally-mediated expression of GCaMP6s under control of the *mDlx* enhancer, previously shown to drive expression specifically in GABAergic cells (Dimidschstein et al., 2016; D. E. Wilson et al., 2017). Wide-field imaging of spontaneous activity was performed at postnatal day 23-29 (P23-29), approximately 7 days before eye opening (typically between P31-34), at a time when modular activity and millimeter-scale networks have been previously shown in excitatory cells (Smith *et al*., 2018) (Figure 3a).

**Figure 3.**
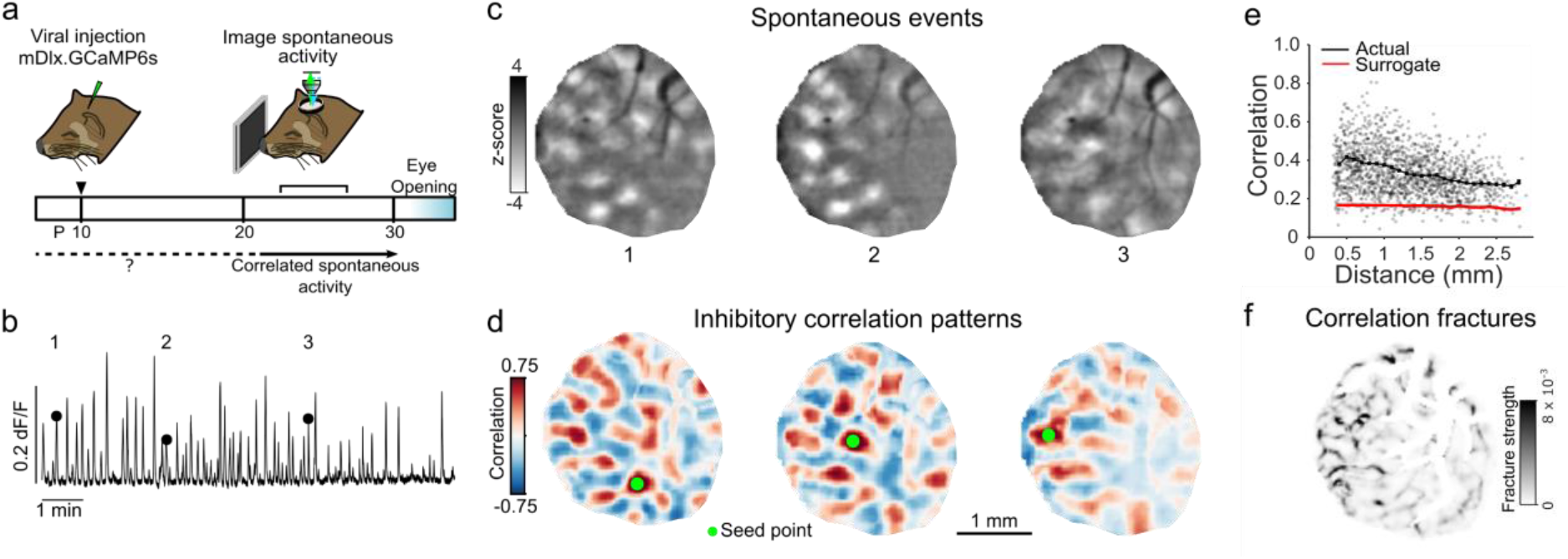
Spontaneous inhibitory activity is modular and participates in precisely organized large-scale correlated networks. **a.** Experimental schematic; ferret kits were injected with inhibitory specific mDlx.GCaMP6s at around P10, and spontaneous activity was imaged prior to eye opening. **b.** Mean trace of inhibitory spontaneous activity across field of view. Numbers correspond to events in (c). **c.** Example single-frame bandpass filtered events. **d.** Spontaneous correlation patterns for three different seed points. **e.** Correlation values at maxima as a function of distance from the seed point, as compared to surrogate shuffled correlations. **f.** Fractures in correlation patterns reveal regions of rapid transition in global correlation structure.

Under light anesthesia, inhibitory neuron populations showed frequent spontaneous activity that exhibited a highly modular structure, with spontaneous events containing multiple distinct domains of activity distributed across the imaging field-of-view and spanning several millimeters (Figure 3b-c). To determine if these modular patterns of activity reflect large-scale correlated inhibitory networks, we calculated the pixelwise Pearson’s correlation coefficient across all detected spontaneous events. We found that inhibitory neurons indeed exhibit a spatially-extended correlation structure, with both strong positively and negatively correlated domains extending several millimeters from the reference seed point (Figure 3d). Peak values of positively correlated domains were statistically significant up to 2 mm away from the seed point (Figure 3e. p<0.01 vs. surrogate data, 6 of 7 individual animals, bootstrap test), demonstrating the large-scale nature of inhibitory networks in the early cortex.

In the mature cortex, orientation preference maps exhibit smooth variation, punctuated by abrupt discontinuities at orientation pinwheels and fractures (Bonhoeffer and Grinvald, 1991; Ohki *et al*., 2005), an organization shared by inhibitory neurons in the mature ferret (Wilson *et al*., 2017). Notably, prior work identified similar discontinuities in the patterns of distributed correlated activity (termed correlation fractures) in excitatory neurons early in development prior to the emergence of orientation maps (Smith *et al*., 2018). To determine if early inhibitory networks exhibit a similar fine-scale organization, we calculated the rate of change in correlation patterns for all pixels within our field-of-view (Supplemental Figure 1). This analysis revealed the presence of spatially discrete fractures over which inhibitory correlation patterns exhibit abrupt changes in structure, demonstrating a precise local organization in the coupling to large scale inhibitory networks in the early visual cortex (Figure 3f).

### Quantitative agreement between inhibitory and excitatory network structure in early cortex

The presence of modular activity patterns, together with long-range correlations exhibiting precise local structure, suggests that inhibitory neurons may already be tightly integrated into functional networks with excitatory cells in the early cortex. To begin to address this, we undertook a quantitative comparison of excitatory and inhibitory networks assessed through spontaneous activity. We performed wide-field imaging in animals expressing either hSyn.GCaMP6s, which is excitatory-specific in ferret (Wilson *et al*., 2017), or the inhibitory-specific mDlx.GCaMP6s. We first quantified the size of modular active domains in spontaneous events by fitting a two-dimensional gaussian ellipse to each active region within an event and calculating the full-width at tenth of maximum (FWTM). Domain size was similar between inhibitory and excitatory events (Figure 4a. minor axis: I: 412.06μm (386.73-439.94), n=7 animals; E: 392.68μm (371.46-397.49), n=7 animals; median (inter-quartile range (IQR)); p=0.142, Wilcoxon rank-sum test), supporting the idea that the modular domains of local inhibitory activity operate on the same scale as excitatory activity.

**Figure 4.**
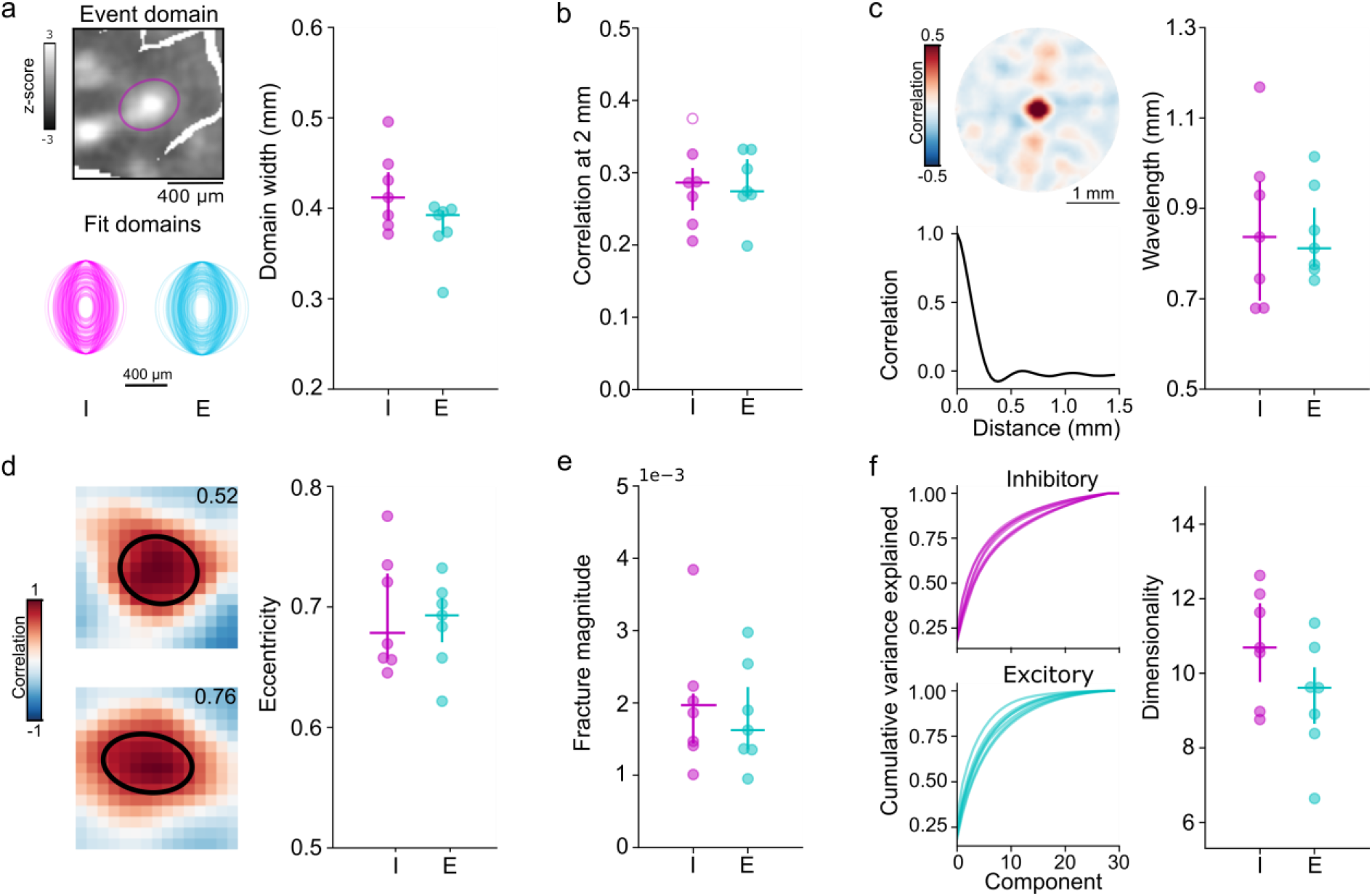
Quantitative agreement in structure of early excitatory and inhibitory networks. **a**. Similar domain size for excitatory and inhibitory events. (Left top) Example event showing FWTM of gaussian fit of active domain. (Left bottom) FWTM for 200 randomly selected domains from individual inhibitory (I) and excitatory (E) animals, rotated to align major axis (Right) Median domain size for each I and E animal. Circles: individual animals, lines: group median and IQR. **b**. Correlation strength at 1.8-2.2 mm from seed point. Filled circles individually significant vs. surrogate (p < 0.01). **c.** Correlation wavelength is similar between I and E networks. (Left top) Correlation pattern averaged over all seed points in single animal with inhibitory GCaMP. (Left bottom) Correlation as function of distance. (Right) Circles: individual animals, lines: group median and IQR. **d.** Similar eccentricity of local correlation structure. (Left) Examples for two seed points from the same inhibitory animal, numbers indicate measured eccentricity. (Right) Circles: individual animals, lines: group median and IQR. **e.** Fracture strength is similar for I and E networks. **f.** Spontaneous activity is moderately low dimensional in both E and I networks. (Left) Cumulative explained variance for principle components of spontaneous events (lines indicate individual animals). (Right) Circles: individual animals, lines: group median and IQR.

Next, we quantitatively examined the correlation patterns for excitatory and inhibitory networks revealed by spontaneous activity. We find that correlations were similarly long-range, with equivalent strength 2 mm away from the seed point (Figure 4b. I: r=0.29 (0.25 - 0.31), E: r=0.24 (0.27 - 0.31); p=0.848, Wilcoxon rank-sum test). Likewise, the wavelength estimated from the full angle-averaged correlation function was similar between inhibitory and excitatory networks (Figure 4c. I: 0.836 mm (0.696 - 0.960), E: 0.811 mm (0.771 - 0.901); p=0.848, Wilcoxon rank-sum test), and approximately similar to the wavelength of orientation columns in the mature cortex (Bonhoeffer and Grinvald, 1993; White, Coppola and Fitzpatrick, 2001; Kaschube *et al*., 2010). Additionally, correlations in inhibitory networks exhibited a strong local anisotropy, with local correlations around the seed point showing a high degree of eccentricity that was consistent with excitatory correlations (Figure 4d. I: 0.699 (0.657 - 0.728), E: 0.693 (0.671 - 0.707); p=0.949, Wilcoxon rank-sum test). Lastly, we found that the strength of correlation fractures was highly similar between excitatory and inhibitory networks, reflecting a similar degree of precision in fine-scale network organization (Figure 4e. I: 1.86*10^−3^ (1.49*10^−3^-2.13*10^−3^), E: 1.62*10^−3^ (1.35*10^−3^-2.22*10^−3^); p=0.655, Wilcoxon rank-sum test). Together, these results demonstrate a strong degree of quantitative similarity in the structure of inhibitory and excitatory networks in the early visual cortex.

Prior work has shown that in locally heterogeneous network models that recapitulate the structure of early excitatory networks, long-range correlations exhibiting both local anisotropy and pronounced fractures strongly coincide with spontaneous activity patterns that reside in a low dimensional subspace (Smith *et al*., 2018). Given the similarities in both local and long-range correlation structure, as well as fracture strength between excitatory and inhibitory networks, we computed the dimensionality of inhibitory spontaneous events and compared them to event-number-matched events in excitatory neurons. We found that across animals, inhibitory and excitatory events tended to reside in similarly low dimensional subspaces (Figure 4f. I: 10.55 (9.60 - 11.78), E: 9.44 (8.44 - 10.01); median (IQR); p=0.142, Wilcoxon rank-sum test), further supporting the contribution of inhibition to local heterogenous networks in the early cortex.

### Precise spatial alignment of inhibitory and excitatory networks across millimeters

These results strongly suggest that excitatory and inhibitory neurons are tightly coupled into the same functional networks early in development. To address this directly, we performed wide-field imaging of spontaneous activity of both excitatory and inhibitory neurons within the same animals (P25-26). Animals were injected with AAVs expressing GCaMP6s in inhibitory cells under control of the *mDlx* enhancer (Dimidschstein *et al*., 2016) and jRCaMP1a (Dana *et al*., 2016) in excitatory cells with hSyn (Wilson *et al*., 2017). Spontaneous activity was imaged using appropriate excitation and emission filters in interleaved blocks of 10 minutes. We observed similar patterns of modular spontaneous events in inhibitory and excitatory neurons, and individual events with corresponding patterns of activity could frequently be found in both excitatory and inhibitory datasets (Figure 5a, Supplemental Figure 2).

**Figure 5.**
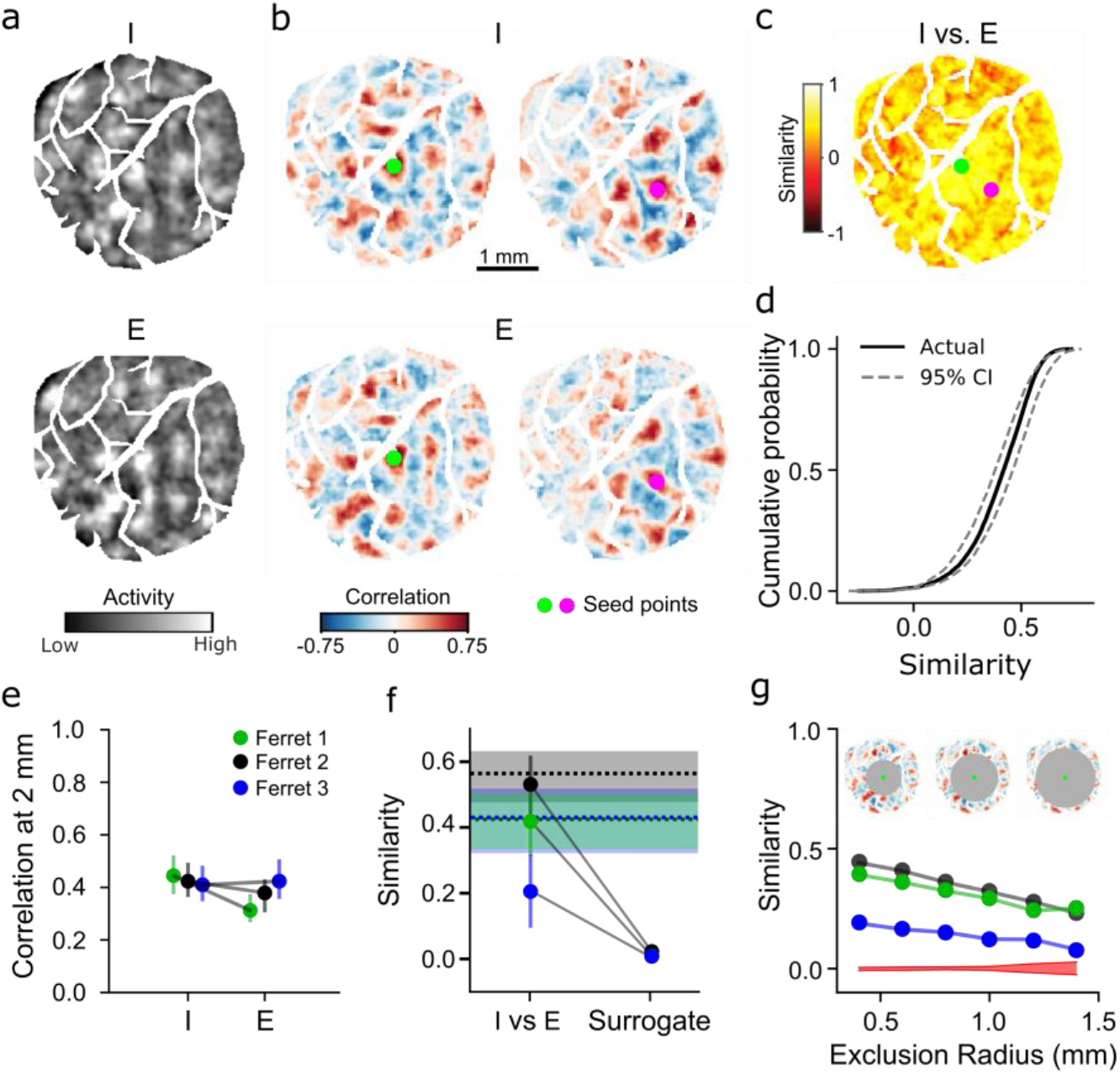
Inhibitory and excitatory networks show high degree of similarity within animal. **a.** Example spontaneous inhibitory (mDlx.GCaMP6s, top) and excitatory (syn.jRCaMP1a, bottom) events recorded from same animal (I color axis: −2-2 z-score, E color axis: −1-1 z score). Example events were chosen based on pattern similarity. **b.** Highly similar correlation patterns in inhibitory and excitatory networks for corresponding seed points. **c.** Quantification of I vs. E correlation similarity for all seed points. Circles indicate seed points illustrated in (b). **d**. Similarity for all seed points in (c) falls within 95% confidence intervals of bootstrapped I vs I similarity. **e.** Within animal comparison of correlation strength for E and I (median and IQR within animal). Filled circles individually significant vs. surrogate (p < 0.01). **f.** Similarity of I vs E correlations is significantly greater than shuffle (median and IQR within animal). Shaded bars indicate IQR of bootstrapped I vs I similarity. **g.** Long-range correlations show significant network similarity. Correlation similarity remains significant vs. surrogate (red, 95% CI of surrogate, averaged across animals) for increasingly distant regions (excluding correlations within 0.4 – 1.4 mm from the seed point).

To compare the structure of excitatory and inhibitory networks, we computed pixelwise correlations separately for all excitatory and inhibitory events for each animal. When selecting the same seed point, the spatial patterns of correlated activity were highly similar across networks (Figure 5b), and exhibited equivalent strength extending across several millimeters (Figure 5e). To quantify this similarity, we computed second-order correlations between excitatory and inhibitory correlation matrices (see methods), which revealed high levels of correlation similarity across nearly all seed-points within the imaging window (Figure 5c), indicating highly similar large-scale networks. To quantify this similarity, we compared it to both the distribution obtained from subsampling inhibitory activity (thereby establishing an upper bound given measurement noise and finite event numbers), as well as to surrogate data. We find that excitatory and inhibitory correlation similarity is significantly greater than surrogate (p<0.01 for 3 of 3 individual animals, bootstrap test) and near the level of within inhibitory similarity (Figure 5d,f; colors correspond to data from 3 individual animals). Notably, excitatory and inhibitory networks exhibit strong similarity even in their long-range correlations, as similarity remained significantly greater than surrogate even when considering only correlations over 1.4 mm from the seed point (Figure 5g. p<0.01, I vs E networks vs surrogate, bootstrap test.) Together, these results demonstrate the presence of overlapping and co-aligned excitatory and inhibitory networks in the early cortex.

### Excitatory and inhibitory networks are tightly integrated at cellular scale

To assess whether the large-scale alignment of excitatory and inhibitory networks observed above extends to the cellular level, we performed simultaneous two-photon imaging of excitatory and inhibitory neurons (P23-26). We co-injected AAVs expressing hSyn-GCaMP6s (excitatory) and mDlx-GCaMP6s-P2A-NLS-tdTomato (inhibitory), allowing us to distinguish inhibitory neurons *in vivo* through tdTomato expression (Figure 6a, Supplemental Figure 3). We observed highly modular spontaneous activity in both excitatory and inhibitory cells, with local populations showing tightly coordinated patterns of activity across cell types (Figure 6b, Supplemental Figure 4).

**Figure 6.**
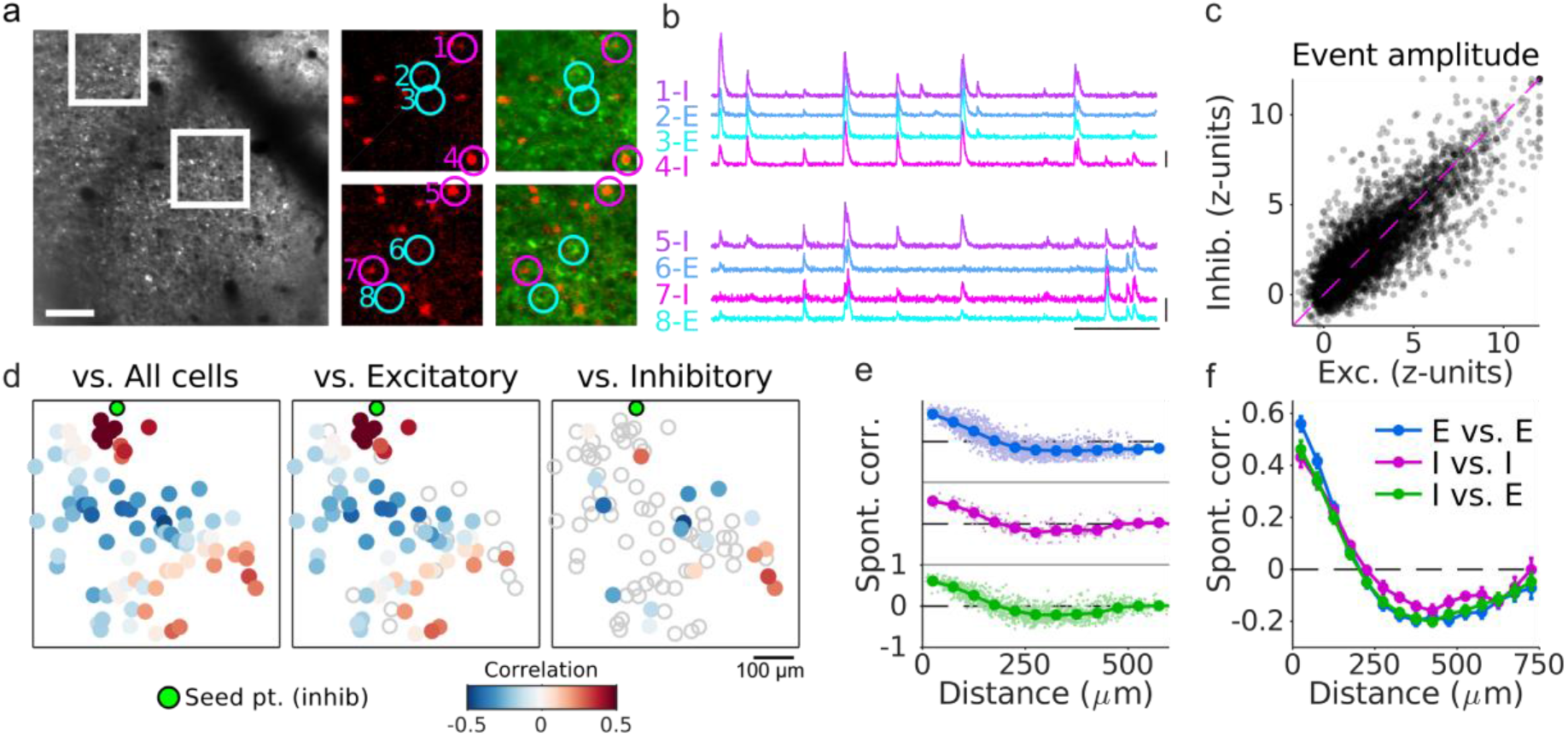
Cellular alignment of excitatory and inhibitory networks in early visual cortex. **a.** (Left): Example FOV showing GCaMP signal from excitatory and inhibitory neurons. (Middle/Right): Expanded images showing nuclear tdTomato signal specifically in inhibitory neurons (middle) and merged with GCaMP signal (right). **b.** Traces of spontaneous activity for excitatory (E, cyan) and inhibitory (I, magenta) neurons as indicated in (a). **c**. Event amplitude for local (<75 μm) excitatory and inhibitory responses. For clarity when plotting, 5% of responses are shown for 2186 cells and 1231 events from 21 FOV in 5 animals, and responses are clipped at 12 z-units. **d**. Example pairwise correlations between an inhibitory neuron (green) and all cells (left), excitatory cells (middle), and inhibitory cells (right). **e**. Pairwise correlations as function of distance for example FOV shown in (ad) for E-E, I-I, and E-I pairs. **f**. Same as (e) averaged over 21 FOV from 5 animals. Scale bars: a,c: 100 μm; b: 30 sec, 3 ΔF/F.

To quantify the relative balance of excitatory and inhibitory activity within local populations, we computed the average response of all cells of a given cell-type within a 75 μm region around each neuron for all spontaneous events. We found the amplitude of events within these local populations was highly correlated between excitatory and inhibitory cells (r=0.86, 2186 cells and 1231 events from 21 FOV in 5 animals; Figure 6c), suggesting locally balanced activity. Both inhibitory and excitatory neurons exhibited spatially organized pairwise correlations across spontaneous events, with distributed patches of positively correlated cells located hundreds of microns away. Notably, the spatial structure of pairwise correlations for a given neuron (whether inhibitory or excitatory) with either inhibitory or excitatory cells was well aligned (p<0.001 vs shuffle), irrespective of cell type (Figure 6d, Supplemental Figure 5). Finally, examining correlation strength as a function of distance and cell type demonstrates that correlated excitatory and inhibitory networks exhibit a similar spatial scale at the cellular level (Figure 6e,f). Together, these results show that excitatory and inhibitory neurons are precisely organized in the early visual cortex into the same spatially structured and locally correlated functional networks.

## Discussion

By applying inhibitory interneuron-specific expression of fluorescent calcium sensors to the early visual cortex, we were able to directly assess the structure of inhibitory networks in the developing cortex and their integration with excitatory networks. We show that prior to eye opening and the onset of reliable stimulus-evoked responses (Chapman, Stryker and Bonhoeffer, 1996), intracortical inhibition is both present and highly organized, with inhibitory networks already displaying patterns of modular and correlated spontaneous activity that span several millimeters. Inhibitory networks exhibit quantitatively similar structure to excitatory networks, which together show precise alignment at both local and global scales in the patterns of correlated spontaneous activity. Together, these findings demonstrate the presence of tightly-coupled excitatory and inhibitory functional networks in the early visual cortex.

In the mature ferret, both inhibitory and excitatory neurons are organized into a columnar map of orientation preference, with orientation-specific subnetworks of functionally coupled excitatory and inhibitory neurons present not only within iso-orientation domains but also near orientation pinwheels (Wilson *et al*., 2017). This organization stands in contrast to the non-specific local pooling of nearby excitatory activity by inhibitory neurons found in the ‘salt-and-pepper’ rodent cortex (Kerlin *et al*., 2010; Bock *et al*., 2011; Hofer *et al*., 2011; Packer and Yuste, 2011; Runyan and Sur, 2013; Scholl *et al*., 2015), raising the possibility that the functionally-specific organization of inhibitory cells in the mature ferret emerges from an initially non-specific state over the course of development. However, our finding of sharp correlation fractures in early inhibitory networks (Figure 3f, Supplemental Figure 1) is inconsistent with non-specific local pooling and argues strongly against this possibility. Rather, our results indicate that in the early cortex, inhibitory neurons are already tightly integrated into functionally-specific networks. Furthermore, given that the patterns of correlated activity undergo extensive refinement in the week prior to eye opening (Smith *et al*., 2018), these results also suggest that excitatory and inhibitory networks likely refine in parallel over development to produce the tightly coupled organization found in the mature cortex (Wilson *et al*., 2017)).

The *mDlx* enhancer used in this study has been shown to drive expression across several subtypes of inhibitory cells (Dimidschstein *et al*., 2016), which have been ascribed distinct roles within cortical circuits (see (Hattori *et al*., 2017; Wood, Blackwell and Geffen, 2017) for reviews). However, in the mature ferret, parvalbumin (PV), somatostatin (SOM) and non-PV, non-SOM GABAergic neurons were all found to be equivalently integrated into specific and spatially-organized functional networks (Wilson *et al*., 2017). Although the low expression of PV in the weeks prior to eye-opening (Gao *et al*., 2000) makes it difficult to separate inhibitory neurons by subtype at these ages, our results suggest that a similarly organized and integrated structure to that in the mature ferret may already exist in the developing cortex. Future work, potentially leveraging inhibitory subtype-specific viral approaches (Mehta *et al*., 2019; Vormstein-Schneider *et al*., 2020), will be required to directly test the functional roles of specific GABAergic subtypes in developing cortical networks.

Large-scale distributed functional networks are a hallmark of mature cortical organization and are exemplified by the columnar arrangement of orientation preference in the visual cortex of primates and carnivores (Blasdel and Salama, 1986; Bonhoeffer and Grinvald, 1991; Ohki *et al*., 2005). Here, co-tuned columns preferentially interconnected by specific long range horizontal projections (Gilbert and Wiesel, 1989; Malach *et al*., 1993; Bosking *et al*., 1997) which emerge over the course of development (Ruthazer and Stryker, 1996; Borrell and Callaway, 2002). The presence of modular activity and long range-correlations in the early cortex prior to the emergence of these horizontal connections, coupled with the finding that correlations persist in the absence of feedforward input (Smith *et al*., 2018), has been taken to indicate that local intracortical circuits are sufficient to generate these patterns of activity. Models of developing cortical networks can rely on local connectivity to self-organize, thereby producing modular patterns of activity (von der Malsburg, 1973; Swindale, 1982; Miller, 1994; Barrow, Bray and Budd, 1996). If these local connections are sufficiently heterogeneous, the resulting patterns of activity reside in a low-dimensional sub-space and produce long range correlations without requiring long-range horizontal connections (Smith *et al*., 2018). A strong prediction of such models is the presence of tightly integrated and co-aligned excitatory and inhibitory networks (Ermentrout and Cowan, 1979), in agreement with our experimental observations in the early cortex. Thus, our results suggest that short-range intracortical interactions between tightly coupled excitatory and inhibitory circuits give rise to large-scale distributed networks during early development, which may serve as a seed for future functional organization in the cortex.

## Acknowledgments

The authors wish to thank Drishti Lall, Casey Xamonthiene, Hailey Glewwe, and Matt Paruzynski for histology and surgical assistance, and members of the Smith and Kaschube labs for helpful discussions. Authors were supported by NIH R01EY030893-01 (GBS), T32 MH115886 (HM), BMBF 01GQ2002 (MK), NSF 1707398 (BH), Gatsby Charitable Foundation GAT3708 (BH), Whitehall Foundation 2018-05-57 (GBS), as well as support from NIH grants P41 EB027061 and P30 NS076408. All viral vectors used in this study were generated by the University of Minnesota Viral Vector and Cloning Core (Minneapolis, MN). This work was supported by the resources and staff at the University of Minnesota University Imaging Centers (SCR 020997).

## Author Contribution

H.M. and G.S. designed the experiments. H.M. and G.S. performed the viral injections and collected the calcium imaging data. H.M. and G.S. analyzed the data, with help from B.H. and M.K. H.M and G.S. wrote the manuscript with input from all authors.

## Declaration of interests

The authors declare no competing interests.

## Materials and Methods

### Animals

All experimental procedures were approved by the University of Minnesota Institutional Animal Care and Use Committee and were performed in accordance with guidelines from the US National Institutes of Health. We obtained 20 male and female ferret kits from Marshall Farms and housed them with jills on a 16-h light/8-h dark cycle. No statistical methods were used to predetermine sample sizes, but our sample sizes are similar to those reported in previous publications.

### Viral Injection

Viral injections were performed as previously described (Smith and Fitzpatrick, 2016). Briefly we expressed GCaMP6s in inhibitory interneurons by microinjecting AAV1-mDLx-GCaMP6s-P2A-NLS-tdTomato (University of Minnesota Viral Vector and Cloning Core), based on the mDlx inhibitory specific enhancer (Dimidschstein *et al*., 2016), into layer 2/3 of visual cortex at P10-P15 approximately 10–15 days before imaging experiments. To image excitatory and inhibitory activity, animals used for widefield experiments were injected with a 1:1 ratio of AAV1-mDlx-GCaMP6s-P2A-NLS-tdTomato and AAV1-Syn-NES-jRCaMP1a-WPRE-SV40, and animals used for two-photon experiments were injected with a 1:1 ratio of AAV1-mDLx-GCaMP6s-P2A-NLS-tdTomato and AAV1-Syn-GCaMP6s-WPRE-SV40.

Anesthesia was induced with isoflurane (3.5-4%) and maintained with isoflurane (1–1.5%). Buprenorphine (0.01mg/kg) and either atropine (0.2 mg/kg) or glycopyrrolate (0.01 mg/kg) were administered, as well as 1:1 lidocaine/bupivacaine at the site of incision. Animal temperature was maintained at approximately 37 °C with a water pump heat therapy pad (Adroit Medical HTP-1500, Parkland Scientific). Animals were also mechanically ventilated and both heart rate and end-tidal CO2 were monitored throughout the surgery. Using aseptic surgical technique, skin and muscle overlying visual cortex were retracted, and a small burr hole was made with a handheld drill (Fordom Electric Co.). Approximately 1 μL of virus contained in a pulled-glass pipette was pressure injected into the cortex at two depths (^~^200 μm and 400 μm below the surface) over 20 min using a Nanoject-II (World Precision Instruments). The craniotomy was filled with 2% agarose and sealed with a thin sterile plastic film to prevent dural adhesion.

### Cranial window surgery

On the day of experimental imaging, ferrets were anesthetized with 3%–4% isoflurane. Atropine was administered as in virus injection procedure. Animals were placed on a feedback-controlled heating pad to maintain an internal temperature of 37 to 38 C. Animals were intubated and ventilated, and isoflurane was delivered between 1 and 2% throughout the surgical procedure to maintain a surgical plane of anesthesia. An intraparietal catheter was placed to deliver fluids. EKG, end-tidal CO2, and internal temperature were continuously monitored during the procedure and subsequent imaging session. The scalp was retracted and a custom titanium headplate adhered to the skull using C&B Metabond (Parkell). A 6 to 7 mm craniotomy was performed at the viral injection site and the dura retracted to reveal the cortex. One 4mm cover glass (round, #1.5 thickness, Electron Microscopy Sciences) was adhered to the bottom of a custom titanium insert and placed onto the brain to gently compress the underlying cortex and dampen biological motion during imaging. The cranial window was hermetically sealed using a stainless-steel retaining ring (5/16-inch internal retaining ring, McMaster-Carr). Upon completion of the surgical procedure, isoflurane was gradually reduced (0.6 to 0.9%) and then vecuronium bromide (0.4 mg/kg/hr) mixed in an LRS 5% Dextrose solution was delivered IP to reduce motion and prevent spontaneous respiration.

### Widefield epifluorescence and two-photon imaging

Widefield epifluorescence imaging was performed with a sCMOS camera (Zyla 5.5, Andor; Prime BSI express, Teledyne) controlled by μManager (Edelstein *et al*., 2010). Images were acquired at 15 Hz with 4 × 4 binning to yield 640 × 540 pixels (Zyla) or 2×2 binning and additional offline 2×2 binning to yield 512 × 512 pixels (Prime BSI). Two-photon imaging was performed with a commercial microscope (Neurolabware) driven by an Insight X3 laser (Spectra Physics). Imaging was performed at 920 nm (GCaMP) and 1040 nm (tdTomato), and fluorescence was collected on separate PMTs using a 562 nm dichroic mirror and 510/84 nm (GCaMP) and 607/70 nm (tdTomato) emission filters (Semrock). Images were collected at 796 × 512 pixels at 30 Hz.

Spontaneous activity was captured in 10-minute imaging sessions, with the animal sitting in a darkened room facing an LCD monitor displaying a black screen.

### Immunostaining and imaging

To confirm that tdTomato expression from AAV1-mDLx-GCaMP6s-P2A-NLS-tdTomato was specific to inhibitory neurons as expected from previous work (Wilson *et al*., 2017), we performed immunostaining in a subset of animals. Following imaging, animals were euthanized and transcardially perfused with 0.9% heparinized saline and 4% paraformaldehyde. The brains were extracted, post-fixed overnight in 4% paraformaldehyde, and stored in 0.1 M phosphate buffer solution. Using a vibratome, brains were tangentially sectioned along the surface of the imaging window (50 μm steps). Slices were stained for GAD67 using Mouse anti-GAD67 (1:1000, Sigma Aldrich, MAB5406) and Alexa 405 donkey anti-mouse (1:500, Abcam, AB175659) as described (Wilson *et al*., 2017). Imaging was performed on a confocal microscope (Nikon C2).

### Bicuculline application

To test whether GABAergic signaling exerted net inhibitory effects at the ages examined in this study, we bath applied the GABA(A) antagonist bicuculline methiodide (BMI) directly to early cortex at P21-23 in animals expressing AAV1-Syn-GCaMP6s-WPRE-SV40. To apply BMI, the cannula covering the cranial window was removed and the exposed cortex was gently flushed with ACSF. After collecting 10 minutes of baseline spontaneous activity, 10 μM BMI in ACSF was bath applied to the cortex, and spontaneous activity was imaged for an additional 10 minutes.

## Data analysis

### Signal extraction for widefield epifluorescence imaging

Image series were motion corrected using rigid alignment and an ROI was manually drawn around the cortical region of GCaMP expression.

Additionally, an ROI mask was manually drawn around blood vessels to remove vessel artefacts. The baseline fluorescence (F0) for each pixel was obtained by applying a rank-order filter to the raw fluorescence trace with a rank 70 (for excitatory data) or 190 (for inhibitory data) and a time-window of 30 s. The rank and time window were chosen such that the baseline faithfully followed the slow trend of the fluorescence activity. The baseline corrected spontaneous activity was calculated as (F − F0)/F0 =Δ F/F0.

### Event detection

Detection of spontaneously active events was performed essentially as described (Smith *et al*., 2018). Briefly, we first determined active pixels on each frame using a pixel-wise threshold set to 3 s.d. above each pixel’s mean value across time. Active pixels not part of a contiguous active region of at least 0.01 mm^2^ were considered ‘inactive’ for the purpose of event detection. Active frames were taken as frames with a spatially extended pattern of activity (>50% of pixels were active). Consecutive active frames were combined into a single event starting with the first high-activity frame and then either ending with the last high-activity frame or, if present, an activity frame defining a local minimum in the fluorescence activity. To assess the spatial pattern of an event, we extracted the maximally active frame for each event, defined as the frame with the highest activity averaged across the ROI. While correlation patterns have been shown to be stable with as low as 10 events (Smith *et al*., 2018), datasets with fewer than 30 detected events were excluded from this study.

Due to differences in signal-to-noise ratio in experiments using jRCaMP1a, we used a pixel-wise threshold of 2 s.d to determine active pixels, and frames were >30% of the pixels were active were considered active frames.

### Spontaneous correlation patterns

Spontaneous correlation patters were calculated as previously described (Smith *et al*., 2018). Briefly, we applied a Gaussian spatial band-pass filter (sigma_low_= 26 μm and sigma_high_ =195 μm) to the maximally active frame in each event and downsampled it to 160 × 135 (128 × 128, Prime BSI) pixels. The resulting patterns, named spontaneous patterns *A* in the following, were used to compute the spontaneous correlation patterns as the pairwise Pearson’s correlation between all locations *x* within the ROI and the seed point *s*:

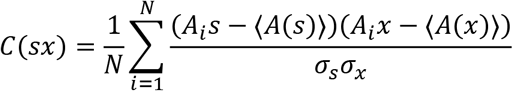

Here the brackets 〈 〉 denote the average over all events and *σx* denotes the standard deviation of *A* over all *N* events *i* at location *x*.

### Spontaneous fractures

Spontaneous fractures were computed as previously described (Smith *et al*., 2018). Fracture strength was defined as the rate by which the correlation pattern changes when moving the seed point location over adjacent pixels. We defined fracture magnitude as the difference in fracture strength averaged over the fracture lines, and its average in regions>130μm apart from the nearest fracture line. Fracture lines were identified by first applying a spatial median filter with a window size of 78μm to remove outliers. We then applied histogram normalization, contrast enhancement using contrast-limited adaptive histogram equalization (CLAHE, clip limit=20, size of neighborhood 260×260 μm^2^), and a spatial high-pass filter (Gaussian filter, s.d. sigma_high_=390 μm). The resulting values were binarized (threshold=0), and the resulting two-dimensional binary array eroded and then dilated (twice) to remove single noncontiguous pixels. We skeletonized this binary array to obtain the fracture lines.

### Event domain size

To estimate the size of active domains in spontaneous events, we first identified domains by taking the local maxima of each event after bandpass filtering. The local neighborhood of each domain (600 μm radius from maxima) was then fit with a two-dimensional gaussian using non-linear least squares:

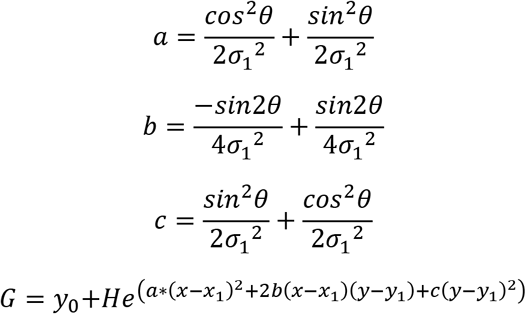

Where *H* is the amplitude of the gaussian, *x*_1_ (*y*_1_) is the *x* (*y*) center, *σ_x_* (*σ_y_*) is the standard deviation of the *x* (*y*) component, *θ* is the angular rotation of the gaussian, and *y_0_* is the offset. Domain size was calculated as the full-width at a tenth of the maxima (FWTM) of the minor axis of fitted gaussian:

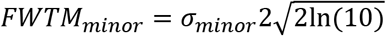

### Strength of long-distance correlations

To determine the strength of correlations, we first identified local maxima (minimum separation between maxima: 800 μm) in the correlation pattern for each seed point. To assess the statistical significance of long-range correlations ^~^2mm from the seed point, we compared the median correlation strength for maxima located 1.8–2.2mm away against a distribution obtained from 100 surrogate correlation patterns. Surrogate correlation patterns control for correlations that arise from finite sampling by eliminating most of the spatial relationship between patterns (Smith *et al*., 2018). Surrogate correlation patterns were generated from spontaneous events that were randomly rotated (rotation angle drawn from a uniform distribution between 0° and 360° with a step size of 10°), translated (shifts drawn from a uniform distribution between±450 μm in increments of 26 μm, independently for x and y directions) and reflected (with probability 0.5, independently at the x and y axes at the center of the ROI).

### Correlation pattern wavelength

To find the wavelength of the correlation patterns, we first centered and averaged the local neighborhood (1500 μm radius) across all seed points. We then collapsed around the angle to obtain the average correlation as a function of distance from the seed point and used spline interpolation to fit the data. The wavelength of the resulting spline interpolation was estimated as the distance to the first local maxima after 0.

### Eccentricity of local correlation structure

To describe the shape of the local correlation pattern around a seed point, we fit an ellipse (least-square fit) with orientation *ϕ*, major axis *ς*_1_ and minor axis *ς*_2_ to the contour line at correlation=0.7 around the seed point. The eccentricity *ε* of the ellipse is defined as:

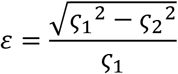

Where ε= 0 is a circle, and increasing values indicating greater elongation along the ellipse.

### Dimensionality of spontaneous activity

We estimated the dimensionality *d*_*ef*f_ of the subspace spanned by spontaneous activity patterns by (Abbott, Rajan and Sompolinsky, 2011):

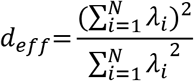

where *λ_i_* are the eigenvalues of the covariance matrix for the N pixels within the ROI. As the value of the dimensionality is sensitive to differences in detected event number, to estimate the distribution of the dimensionality for each animal we calculated the dimensionality of randomly sub-sampled events (n=30 events, matched across animals, 100 simulations) and took the median of the distribution.

### Comparison of inhibitory and excitatory correlation similarity

To compare the similarity between inhibitory and excitatory correlation patterns within the same animal, we computed the second-order correlation between patterns. For each seed point, we calculated the second order Pearson’s correlation between corresponding correlation patterns, while excluding pixels within a 400 μm exclusion radius around the seed point to prevent local correlations from inflating the similarity between the two networks. To get an estimate the upper bound of similarity within inhibitory networks, given a finite sampling size, we randomly split the detected inhibitory events into two groups and separately computed correlations and the second-order correlations between the halves (n simulations=100.) To determine if the observed networks are more similar than chance, we calculated the similarity between the excitatory network and a surrogate inhibitory network (surrogate events calculated as above, with 100 simulations and surrogate similarity calculated as median and IQR across simulations). To determine if spontaneous correlations far from the seed point also maintain high degrees of similarity, we systematically increased the size of the exclusion radii, calculating similarity only using data far from the seed points. Exclusion radius size ranged from 400 to 1400 μm, in 200 μm steps.

### Two-photon event detection and cellular correlations

Two-photon images were corrected for rigid in plane motion via a 2D cross-correlation. Cellular ROIs were drawn using custom software (Cell Magic Wand (Wilson *et al*., 2017)) in ImageJ and imported into Matlab via MIJ (Sage, D., Prodanov, D., Tinevez, J. and Schindelin, J. MIJ: making interoperability between ImageJ and Matlab possible, ImageJ User & Developer Conference, 24–26 October 2012, Luxembourg, http://bigwww.epfl.ch/sage/soft/mij/). Cellular ROIs were then manually categorized as either being inhibitory or excitatory based on tdTomato expression. Fluorescence was averaged over all pixels in the ROI and traces were neuropil subtracted, where F_neuropil_ was taken as the average fluorescence within 30 μm excluding all cellular ROIs and α=0.6:

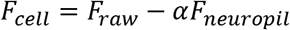

Activity was taken as ΔF/F0, where F0 was the baseline fluorescence obtained by applying a 60-s median filter, followed by a first order Butterworth high-pass filter with a cutoff time of 60s. To compute spontaneous correlations, we first identified frames containing spontaneous events, which were defined as frames in which > 10% of imaged neurons exhibited activity > 2 s.d. above their mean. Cellular activity on all event frames was then z-scored using the mean and s.d. of each frame, and pairwise Pearson’s correlations were computed across all neurons over all active frames.

The similarity of cellular correlation patterns for inhibitory and excitatory cells was computed by first creating a spatially matched set of excitatory and inhibitory neurons. For each inhibitory cell in the field of view, a spatially matched excitatory cell was identified by finding the closest excitatory cell (within 50 μm). Then, for every cell in the field of view (excitatory and inhibitory) pairwise correlations were calculated with either inhibitory cells or the spatially matched excitatory cells. The similarity of these correlation patterns was taken as the second order Pearson’s correlation as above, while excluding cells <200 μm from the seed neuron. Statistical significance was computed by comparing measured similarity values against a shuffled distribution obtained by shuffling the pairwise correlations before computing similarity (100 shuffles).

To compare the amplitude of events within a local area, activity traces for each cell were smoothed with a 3-sample median filter and z-scored across all frames. We then computed the average amplitude of all inhibitory (or excitatory) cells within a 75 μm radius for the maximally active frame of each event.

### Quantification and statistical analysis

Nonparametric tests were used for statistical testing throughout the study. Bootstrapping was used determine null distributions when indicated. Center and spread values are reported as median and inter-quartile range, unless otherwise noted. Statistical analyses were performed in MATLAB and Python, and significance was defined as p < 0.05.

### Data availability

Source data and code (Python, MATLAB) used for analysis in this work are accessible at https://github.com/mulho042/SpontaneousInhib.git. Additional data available upon reasonable request.

## Supplemental Figures

**Supplemental Figure 1.**
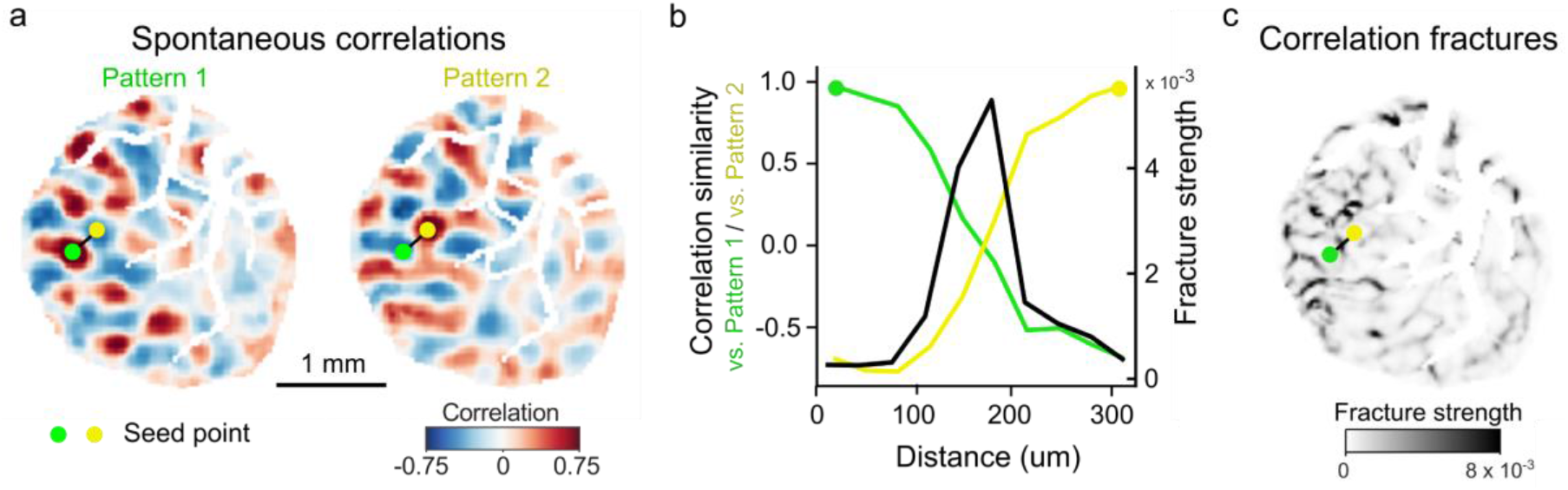
Correlation patterns of inhibitory spontaneous activity exhibit pronounced fractures. **a.** Example correlation patterns showing rapid change in correlation structure between nearby seed points. **b**. Similarity of correlations falls off rapidly as the seed point is moved along the black line in (a), reflected in a sharp peak in fracture strength. **c**. Spatial distribution of fracture strength for all pixels in FOV.

**Supplemental Figure 2.**
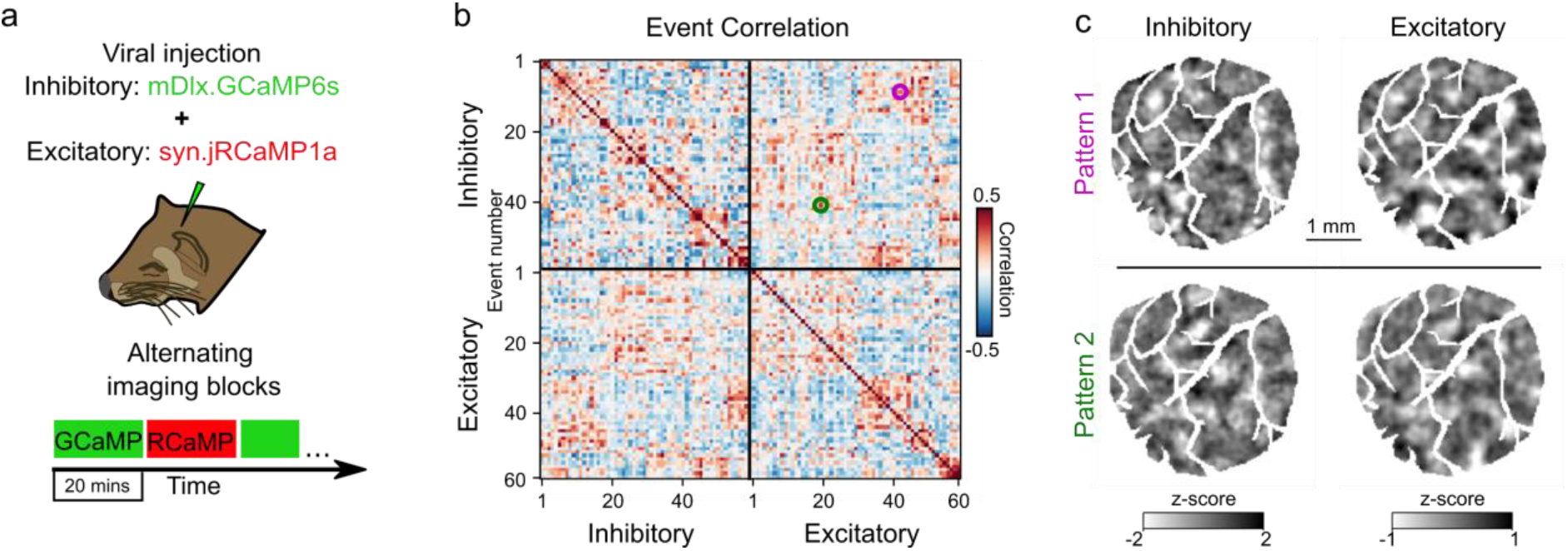
Similar patterns of spontaneous activity in inhibitory and excitatory events. **a.** Experimental schematic; I and E spontaneous activity imaged in interleaved 20 min blocks **b.** Sorted correlation matrix of inhibitory and excitatory events. Matrix shows not only clear clusters of repeating patterns within inhibitory and excitatory events, but also correlated patterns between inhibitory and excitatory events. **c.** Example inhibitory and excitatory events with correlated spatial patterns.

**Supplemental Figure 3.**
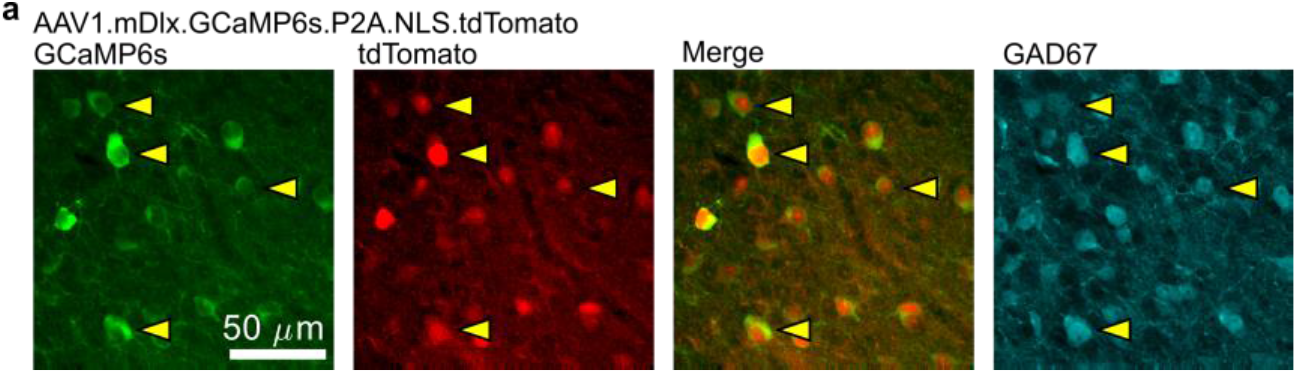
tdTomato expressed from AAV1-mDlx-GCaMP6s-P2A-NLS-tdTomato specifically labels GABAergic neurons. *(From left to right)* Confocal images of GCaMP6s (green), tdTomato (red), merged GCaMP6s and tdTomato, and glutamate decarboxylase 67 (GAD67, cyan). Arrows indicate individual cell bodies, showing co-labeling of GCaMP6s, nuclear tdTomato, and GAD67.

**Supplemental Figure 4.**
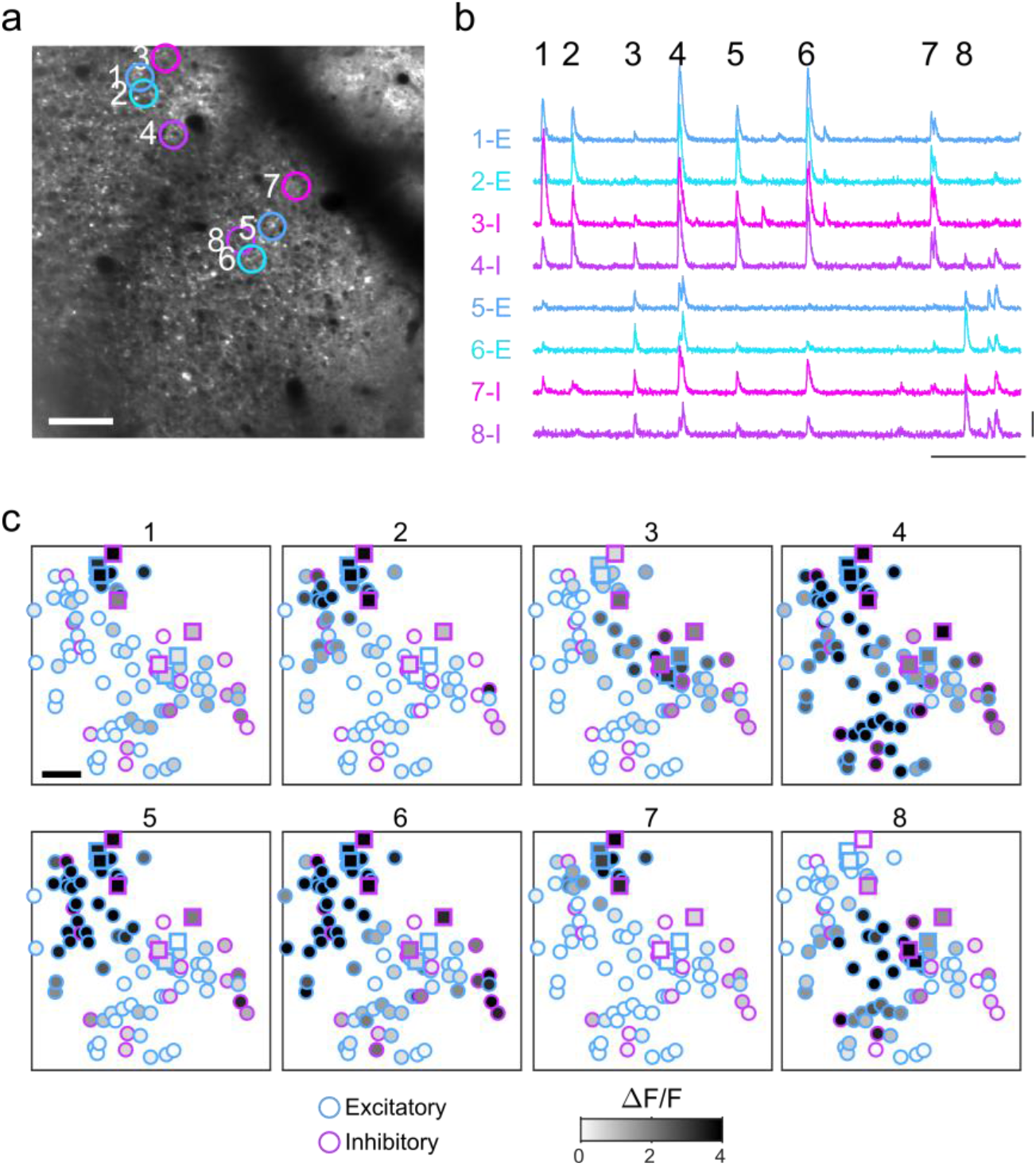
Spontaneous activity patterns are spatially modular and aligned across excitatory and inhibitory neurons. **a**, **b**. Example FOV and activity traces for excitatory and inhibitory neurons shown in (a) (replotted from Figure 4). **c**. Cellular activity patterns for 8 spontaneous events indicated in (b). Neurons shown in (a) & (b) are indicated with square symbols. Scale bars: a,c: 100 μm; b: 30 sec, 3 ΔF/F.

**Supplemental Figure 5.**
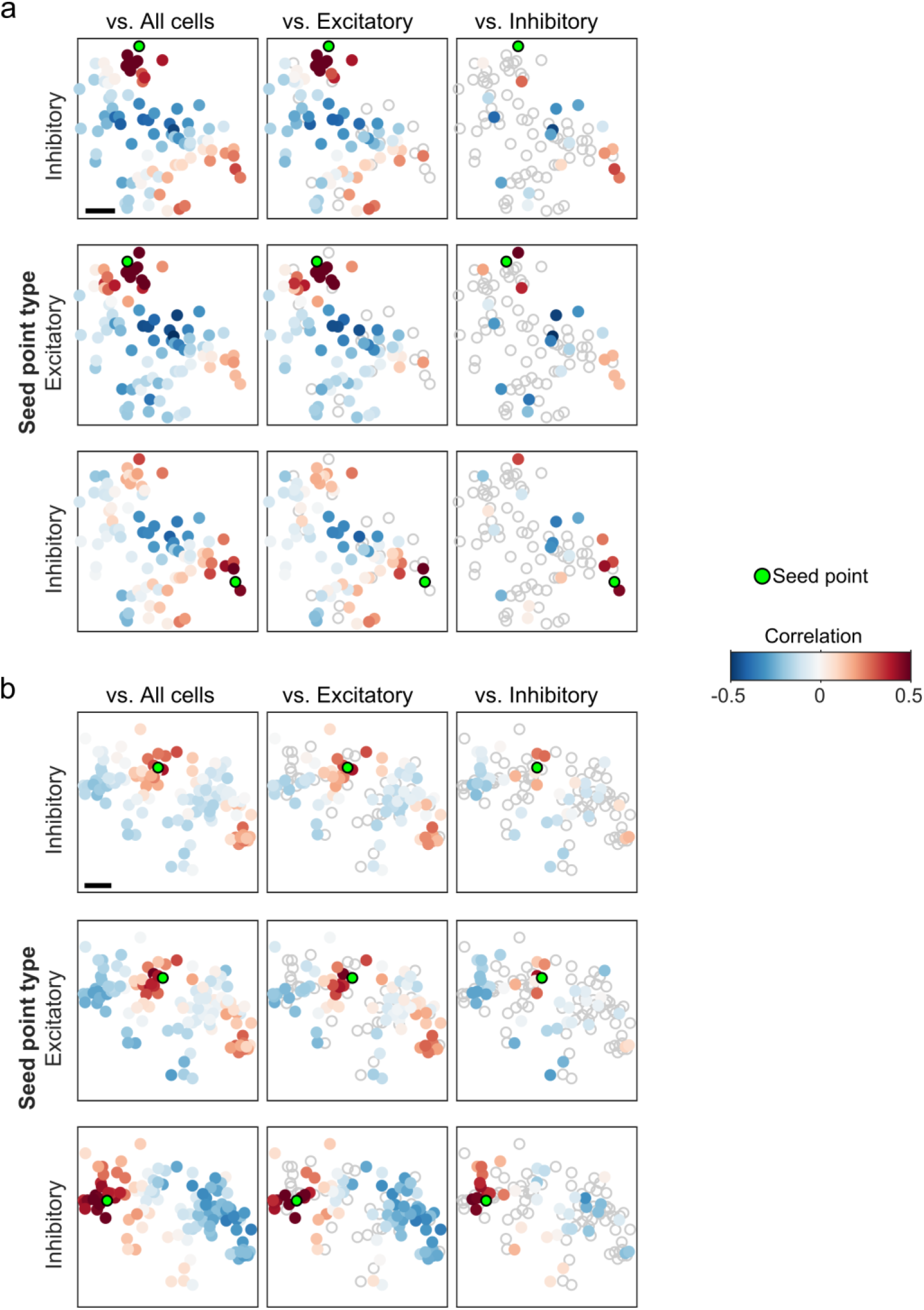
Examples of co-aligned correlation structure between excitatory and inhibitory cells. **a**. Top row: Pairwise correlations between an example inhibitory neuron and all cells (*left*), excitatory cells (*middle*), and inhibitory cells (*right*), replotted from figure 4d. *Middle and bottom* row: additional example seed points (*middle*: excitatory neuron, *bottom*: inhibitory neuron) from same FOV. **b**. Same as (a), but for different animal. Scale bars: 100 μm.

## Notes

### Competing Interest Statement

The authors have declared no competing interest.

